# The DOE Systems Biology Knowledgebase (KBase)

**DOI:** 10.1101/096354

**Authors:** Adam P Arkin, Rick L Stevens, Robert W Cottingham, Sergei Maslov, Christopher S Henry, Paramvir Dehal, Doreen Ware, Fernando Perez, Nomi L Harris, Shane Canon, Michael W Sneddon, Matthew L Henderson, William J Riehl, Dan Gunter, Dan Murphy-Olson, Stephen Chan, Roy T Kamimura, Thomas S Brettin, Folker Meyer, Dylan Chivian, David J Weston, Elizabeth M Glass, Brian H Davison, Sunita Kumari, Benjamin H Allen, Jason Baumohl, Aaron A Best, Ben Bowen, Steven E Brenner, Christopher C Bun, John-Marc Chandonia, Jer-Ming Chia, Ric Colasanti, Neal Conrad, James J Davis, Matthew DeJongh, Scott Devoid, Emily Dietrich, Meghan M Drake, Inna Dubchak, Janaka N Edirisinghe, Gang Fang, José P Faria, Paul M Frybarger, Wolfgang Gerlach, Mark Gerstein, James Gurtowski, Holly L Haun, Fei He, Rashmi Jain, Marcin P Joachimiak, Kevin P Keegan, Shinnosuke Kondo, Vivek Kumar, Miriam L Land, Marissa Mills, Pavel Novichkov, Taeyun Oh, Gary J Olsen, Bob Olson, Bruce Parrello, Shiran Pasternak, Erik Pearson, Sarah S Poon, Gavin A Price, Srividya Ramakrishnan, Priya Ranjan, Pamela C Ronald, Michael C Schatz, Samuel M D Seaver, Maulik Shukla, Roman A Sutormin, Mustafa H Syed, James Thomason, Nathan L Tintle, Daifeng Wang, Fangfang Xia, Hyunseung Yoo, Shinjae Yoo

## Abstract

The U.S. Department of Energy Systems Biology Knowledgebase (KBase) is an open-source software and data platform designed to meet the grand challenge of systems biology — predicting and designing biological function from the biomolecular (small scale) to the ecological (large scale). KBase is available for anyone to use, and enables researchers to collaboratively generate, test, compare, and share hypotheses about biological functions; perform large-scale analyses on scalable computing infrastructure; and combine experimental evidence and conclusions that lead to accurate models of plant and microbial physiology and community dynamics. The KBase platform has (1) extensible analytical capabilities that currently include genome assembly, annotation, ontology assignment, comparative genomics, transcriptomics, and metabolic modeling; (2) a web-browser-based user interface that supports building, sharing, and publishing reproducible and well-annotated analyses with integrated data; (3) access to extensive computational resources; and (4) a software development kit allowing the community to add functionality to the system.

## Introduction

Over the past two decades, the scale and complexity of genomics technologies and data have advanced from simple genomic sequences of only a few organisms to metagenomes and genome variation, gene expression, metabolite, and phenotype data for thousands of organisms and their communities. A major challenge in this data-rich age of biology is integrating heterogeneous, distributed, and error-prone primary and derived data into predictive models of biological function ranging from a single gene to entire organisms and their ecologies. To develop models of these biological processes, organisms and their interactions, scientists of diverse backgrounds need, at a minimum, the ability to use a variety of sophisticated computational tools to analyze their own complex and heterogeneous data sets, and then integrate their data and results effectively with the work of others. Such integration requires discovering and accessing others’ information, understanding its source and limitations, and using tools to perform additional analyses upon it. Ideally, new data and conclusions would be rapidly propagated across existing, related analyses and easily discovered by the community for evaluation and comparison with previous results^1–3^.

Nowhere are the barriers to discovery, characterization, and prediction more formidable than in efforts to understand the complex interplay between biological and abiotic processes that influence soil, water, and climate dynamics and impact the productivity of our biosphere. A bewildering diversity of plants, microbes, animals, and their interactions needs to be discovered and characterized to mechanistically understand ecological function and thereby facilitate interventions to improve outcomes. The U.S. Department of Energy (DOE) has invested in various large- and small-scale programs in climate and environmental science and biological system science. These efforts have demonstrated the power of integrated science programs to make progress on complex systems. However, the community that has grown around these efforts has recognized the need to lower the barrier to accessing tools, data, and results, and to work collaboratively to accelerate the pace of their research^4^.

The DOE Systems Biology Knowledgebase (KBase, www.kbase.us) is a software platform designed to provide these needed capabilities. Specifically, KBase seeks to make it easier for scientists to create, execute, collaborate on, and share sophisticated reproducible analyses of their own biological data in the context of public and other users’ data. Results and conclusions can be shared with individuals or published within KBase’s integrated data model that will increasingly support user-driven and automated meta-analysis.

While a number of recent computational environments address different aspects of this challenge (see *Comparison with other Platforms*), they are generally decentralized, limiting the extent to which data and workflows can be integrated, shared, and extended across the scientific community. Moreover, there is minimal support for key capabilities such as more iterative scientific analysis, in-depth collaboration, integration of new results in the context of others’ results, and automatic propagation of new findings that may inform research across disciplines.

KBase users have already applied the system to address a range of scientific problems, including comparative genomics of plants, prediction of microbiome interactions, and deep metabolic modeling of environmental and engineered microbes. KBase currently supports a growing and extensible set of applications or “apps” for genome assembly, annotation, metabolic model reconstruction, flux balance analysis, expression analysis, and comparative genomics. In addition to these tools, the KBase platform provides data integration and search, along with easy access to shared user analyses of public plant and microbial reference data from a number of external resources including National Center for Biotechnology Information (NCBI) and the DOE Joint Genome Institute (see *KBase Data Model and Apps*). As the platform matures and is adopted more widely, the data, analysis tools, and computational experiments contributed by users are also expected to increase, leading to wider biological applications with richer and more sophisticated support for functional prediction and comparison.

Based on a service-oriented design, KBase is built to run primarily on a cloud-computing infrastructure, although high-performance computing (HPC) resources are only now being integrated. The platform is completely open source, with all core infrastructure and service code available on GitHub (https://github.com/kbase). The central instance of KBase that serves analysis and modeling of plants, microbes, and their communities is maintained and run on DOE enterprise computing resources. This resource is open and free for anyone to use.

### KBase Narratives and User Interface

KBase’s graphical user interface, the Narrative Interface, supports both point-and-click and scripting access to system functionality in a “notebook” environment, enabling computational sophisticates and experimentalists to easily collaborate within the same platform. Built on the Jupyter Notebook^5^, the interface allows researchers to design, carry out, record and share computational experiments in the form of *Narratives*—dynamic, interactive documents that include all the data, analysis steps, parameters, visualizations, scripts, commentary, results, and conclusions of an experiment (Fig. 1).

**Figure 1.**
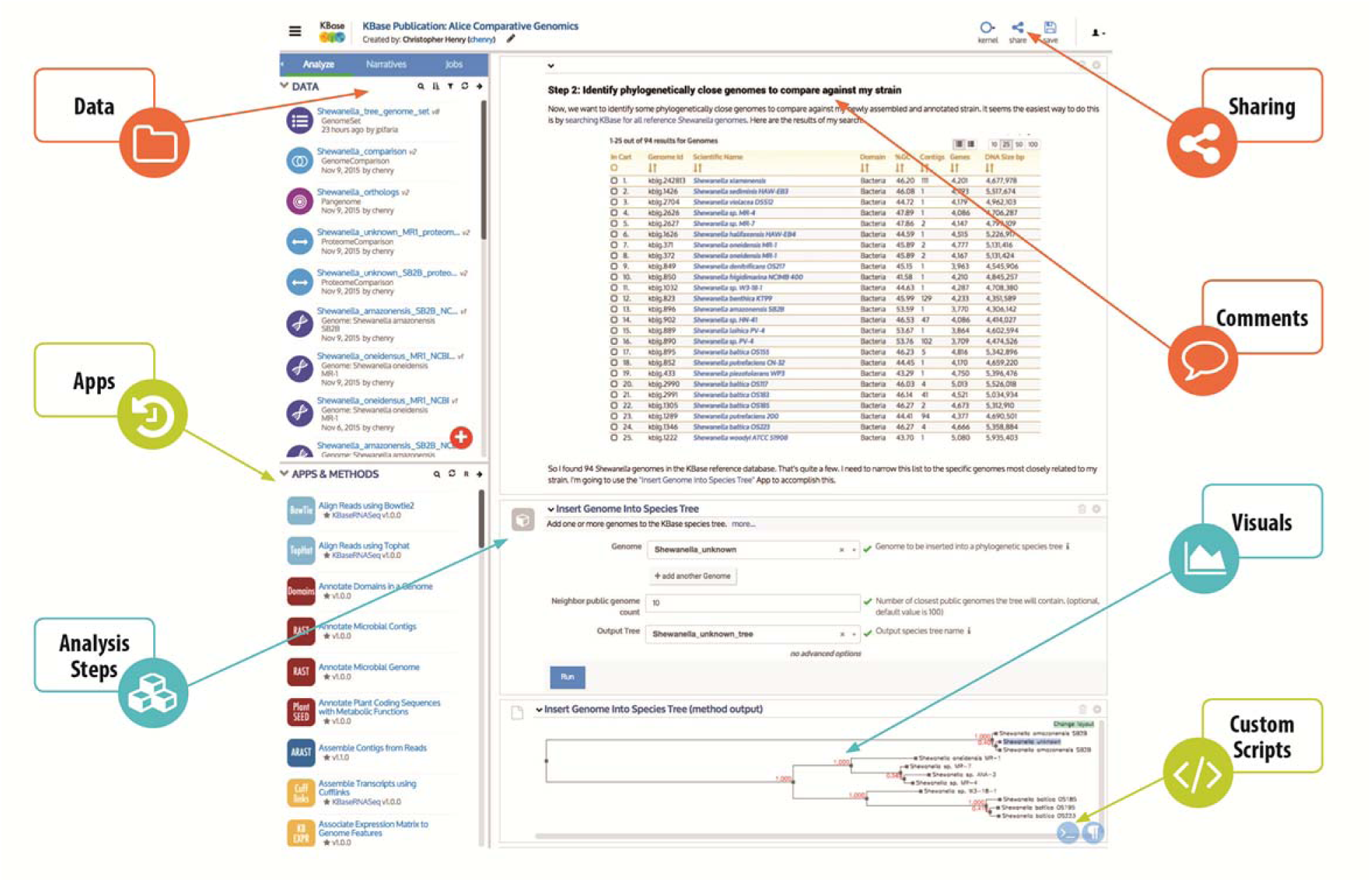
*KBase Narratives. A Narrative is an interactive, dynamic, and persistent document created by users that promotes open, reproducible, and collaborative science*.

Although private by default, users can choose to share their Narratives and data with select collaborators or even publicly. Sharing among collaborators facilitates communication, reuse and management of scientific projects. Public Narratives serve as critical resources for the broader user community by capturing valuable data sets, associated computational analyses, and enriched scientific context describing the rationale and design of original experiments, data upload and organization, and the application of various analytical techniques, complete with the selected parameters and interpretation of results. Narrative sharing and publication are key capabilities enabling other scientists to quickly view, copy, replicate, and expand on work performed within KBase. A growing number of public Narratives are available in KBase today, several of which are described in detail in the “Science Performed within the KBase Platform” section. A selection of these is also available in the KBase Narrative Library (www.kbase.us/narrative-library).

Because Narratives are built upon the Jupyter Notebook framework, users can create and run scripts within a Narrative using a “code cell.” KBase is building a code cell application programming interface (API) enabling users to run KBase apps programmatically from within the system. Users also can leverage the flexibility of code cells to incorporate custom analysis steps into their Narratives not yet available as KBase apps.

### KBase Data Model and Apps

KBase provides a seamless amalgamation of (1) curated and periodically updated reference data, (2) uploaded private and shared user data (and derived data products), and (3) a growing compendium of apps covering a wide range of analytical capabilities. KBase’s reference database includes all public genome sequences from RefSeq^6^ and Phytozome^7^ and selected plant genomes from Ensembl^8^. These genomes are maintained with their original gene IDs and annotations, along with updated gene calls and annotations provided by the KBase annotation pipeline. The reference collection also contains biochemistry data, including 27,692 biochemical compounds, 34,705 reactions, and 529 growth media formulations, and integrates many prominent ontologies such as GO^9^, SEED Subsystems^10^, PFam^11^, and Interpro^12^. All reference data are periodically updated from source databases and are available for integration with user data where appropriate (e.g., when identifying isofunctional genes or building species trees).

KBase’s faceted search utility enables users to query reference data by text, DNA sequence, or protein sequence. Genomes and genes identified through searches can be copied to a Narrative for deeper analysis. Ultimately, the search utility will be extended to query the user data in KBase, while still ensuring appropriate data privacy. This capability will facilitate the process by which users search their own data and data shared with them by collaborators.

In addition to reference data, KBase stores a wide range of uploaded and derived data sets shared by users and connected to analyses performed in Narratives (Fig. 2A). Currently supported data objects include reads, contigs, genomes, metabolic models, growth media, RNA-seq, expression, growth phenotype data, and flux balance analysis solutions. This set of data objects is fully extensible, and the links among them are expanded as new data sources and apps are added to the platform. Third-party developers cannot yet extend the KBase data model, but providing this capability is an important near-term goal.

KBase currently has over 70 released apps in production offering diverse scientific functionality for genome assembly, genome annotation^13^, sequence homology analysis, tree building^14^, comparative genomics, metabolic modeling^15^, community modeling^16^, gap-filling^17,18^, RNA-seq processing^19^, and expression analysis^20^. In addition there are dozens of beta (pre-release) apps available to try. Apps are engineered to interoperate seamlessly to enable a range of scientific workflows (Fig. 2B). Current apps, production and beta, with their associated documentation are listed in the KBase App Catalog (https://narrative.kbase.us/#appcatalog). KBase enables third-party developers to add their own apps for use by the broad scientific community (see KBase Software Development Kit section).

**Figure 2.**
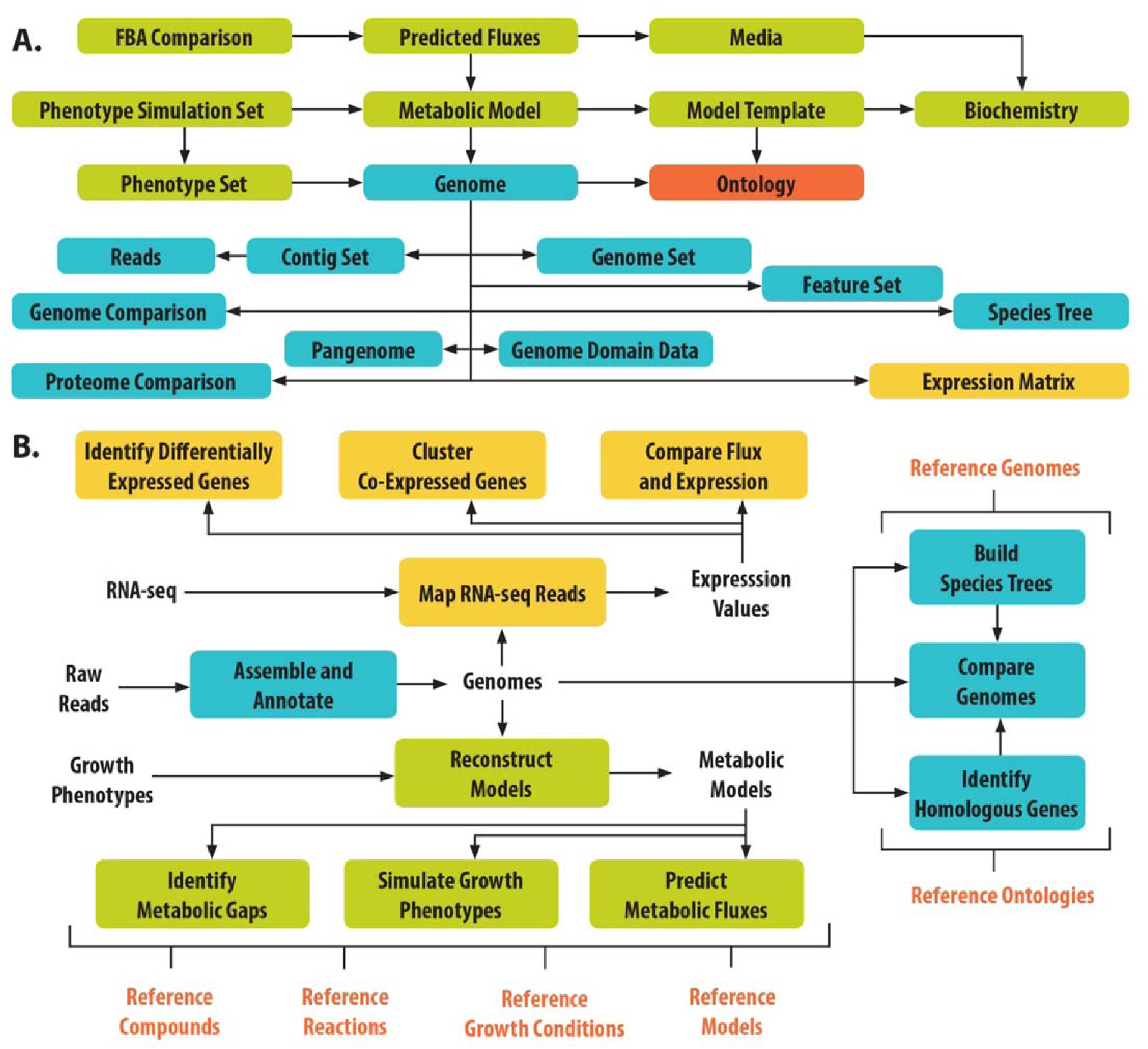
*Currently supported (A) data types and (B) workflows in the KBase platform. KBase integrates data types and apps for analyzing and comparing genomes (blue), transcriptomes (orange), and metabolic models (green)*.

### Support for Reproducible, Interdisciplinary, Collaborative Science

The KBase platform is designed from the ground up to comprehensively support reproducible, interdisciplinary, and collaborative science. KBase’s object-based data store, for example, holds all user and reference data, and all objects, including Narratives themselves, have their own provenance and version information. Provenance data capture when, how, and by whom each object was created, including the apps or upload tools used and their associated parameters and files. All apps and data are also versioned, so that even if a user overwrites an object or an app is updated, the original information is recoverable. These provenance and reversibility features ensure that all science performed within KBase is fully specified and reproducible, averting the time-consuming effort often required to understand how published computational studies or even simple data analyses were carried out.

Collaboration in KBase is supported by the ease with which all data in the system may be shared and copied among users. A user may share any Narrative that they own (or have administrative access to) with other KBase users simply by searching for their names or email addresses. Importantly, when a user shares a Narrative, they also are sharing all the data objects loaded, used, or generated within the Narrative, complete with versioning and provenance. This feature allows users to share every aspect of the science that they have done in KBase. Three levels of sharing are supported: (1) *read* sharing lets others see and copy, but not edit the Narrative and its associated data; (2) *write* sharing expands these privileges to include editing, enabling users to work together on a single Narrative; and (3) *admin* sharing allows others to edit the Narrative and also share it with third parties. Moreover, any Narrative can be made public by the user who owns it, which essentially gives all KBase users read access to that Narrative and its underlying data. Users with read privileges for a Narrative can create their own copy, which they own and can edit. This copying feature enables users to quickly replicate and expand on any KBase Narrative shared with them. This approach to sharing facilitates interdisciplinary science by allowing researchers with different expertise to quickly and easily exchange data, results, methodologies, and workflows used to solve complex biological problems.

We demonstrate how KBase facilitates sharing, collaboration, and interdisciplinary research with a series of example Narratives (Fig. 3 and Table 1) from two scientists: Alice, a wet-lab biologist with expertise in assembly, annotation, and comparative genomics, and Bob, a computational biologist with expertise in metabolic modeling. (Before proceeding to view these Narratives, sign up for a KBase User Account at www.kbase.us) In the first Narrative (Table 1, Narrative 1), Alice uploads raw reads from a new strain of *Shewanella* that she is analyzing. She uses KBase to assemble and annotate these reads, generating a new genome object in KBase. In a second Narrative (Table 1, Narrative 2), Alice compares her new genome with other close strains of *Shewanella*. She finds growth phenotype data for *Shewanella oneidensis* MR-1, which is phylogenetically close to her strain^21^. This inspires Alice to run a growth phenotype array on her own strain, which she also uploads to KBase. Alice then compares both phenotype arrays and notices many differences that she cannot explain.

**Figure 3.**
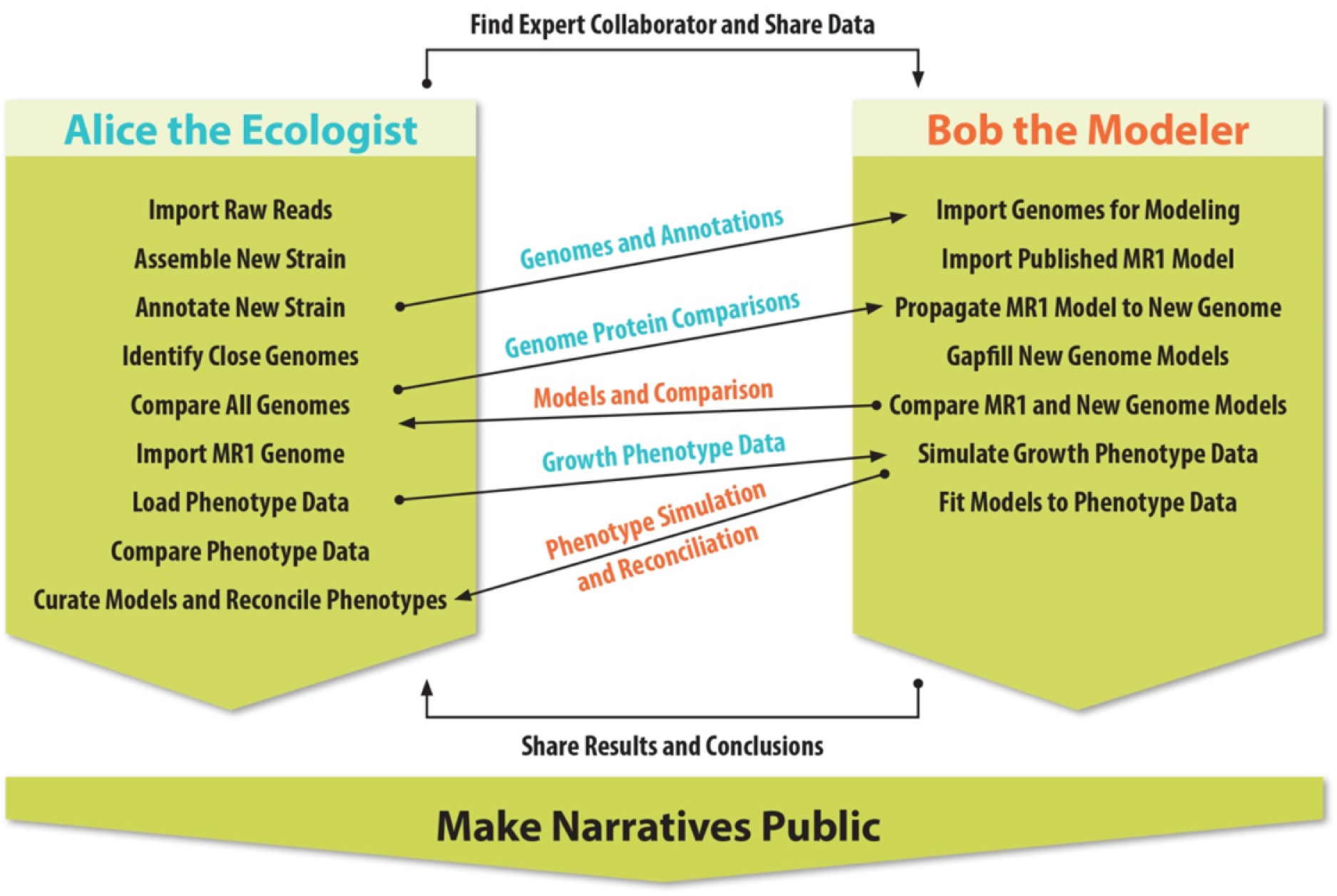
*Example of collaboration in KBase. Two researchers collaborate using Narratives to reach more complete scientific conclusions than either could have achieved alone*.

**Table 1.** Example workflows demonstrating collaboration using KBase. (Sign up for a KBase User Account at www.kbase.us before clicking the links to view these Narratives. This table with active links is also available online at www.kbase.us/kbase-paper.)

| Narrative # | Title and URL |
| --- | --- |
| 1 | Alice Narrative 1: Assembly and Annotation<br>(https://narrative.kbase.us/narrative/ws.18152.obj.1) |
| 2 | Alice Narrative 2: Comparative Genomics<br>(https://narrative.kbase.us/narrative/ws.18153.obj.1) |
| 3 | Bob Narrative: Build Metabolic Models<br>(https://narrative.kbase.us/narrative/ws.18155.obj.1) |
| 4 | Bob and Alice Narrative 1: Phenotype Data Analysis<br>(https://narrative.kbase.us/narrative/ws.18156.obj.1) |
| 5 | Bob and Alice Narrative 2: Phenotype Data Reconciliation<br>(https://narrative.kbase.us/narrative/ws.18157.obj.1) |

At this point, Alice contacts Bob, who suggests that metabolic models can help her analyze the Biolog phenotype data. Alice shares her Narratives with Bob, who copies her genomes into a new third Narrative. In this third Narrative (Table 1, Narrative 3), Bob loads a published model of *S. oneidensis* MR-1^22^, which he then propagates to Alice’s genome. Bob compares the models, identifying some interesting metabolic differences. Then Bob creates a fourth Narrative (Table 1, Narrative 4), where he imports Alice’s Biolog data and simulates the data with his *Shewanella* models. He optimizes his models to fit the Biolog data and shares the results with Alice.

Finally, they build a fifth Narrative (Table 1, Narrative 5) together in which Alice refines Bob’s models by replacing gap-filled reactions with more biologically relevant selections, gaining a complete understanding of the differences between her strain and MR-1. All data used in this example are real: Alice’s raw reads are from an existing genome, *Shewanella amazonensis* SB2B^23,24^, and the growth phenotype data are from an existing experimental study^25^. The computational experiment carried out in the five Narratives results in the development and validation of a new genome-scale metabolic model of *S. amazonensis* SB2B in KBase, as well as the improvement of the existing model for *S. oneidensis* MR-1.

This example demonstrates the novel, collaborative science that can be done in KBase using the extensive portfolio of tools and data currently in the system. KBase provides some new analysis tools (e.g., community modeling^16^ and growth phenotype gap-filling) that are not yet available in any other platform. KBase also makes accessible a number of third-party, open-source tools with the goal of making them easier to use and more powerful when integrated with existing tools in the broader KBase ecosystem. Equally important in this example is how KBase’s user interface facilitates a seamless collaboration between scientists with different but complementary expertise who are able to accomplish more together than they could individually. Finally, it is important to note how KBase reference data (such as genomes, media conditions, and biochemistry) can be integrated into users’ analyses, enhancing their work and offering new perspectives.

### Science Performed Within the KBase Platform

KBase launched the first version of the Narrative user interface in February 2015. Since then, more than 1200 users have signed up for KBase accounts, and over 750 have created at least one Narrative. Excluding KBase staff, users have built 2657 Narratives, 294 of which have been shared with at least one other user and 37 have been made public. These Narratives contain 255,148 data objects, or an average of 96 data objects and five apps per Narrative. In these Narratives, users have applied KBase to address a wide range of scientific questions. Here we highlight seven peer-reviewed publications that cite publicly available Narratives in KBase, where the bulk of the analysis described in the publication was performed. These and other science Narratives can be found in KBase’s Narrative Library, www.kbase.us/narrative-library.

Major applications for KBase include reconstructing, comparing, and analyzing metabolic models for diverse genomes. In 2015, the PlantSEED metabolic model reconstruction pipeline was integrated into KBase and used to build genome-scale metabolic models of 10 diverse reference plant genomes, which were subsequently compared (Table 2, Narrative 1)^26^. More recently, KBase implemented a core model reconstruction pipeline that was applied to more than 8000 microbial genomes (Table 2, Narrative 2)^27^. The resulting core models were compared within a phylogenetic context, identifying how the pathways comprising core metabolism are clustered across the microbial tree of life.

**Table 2.**
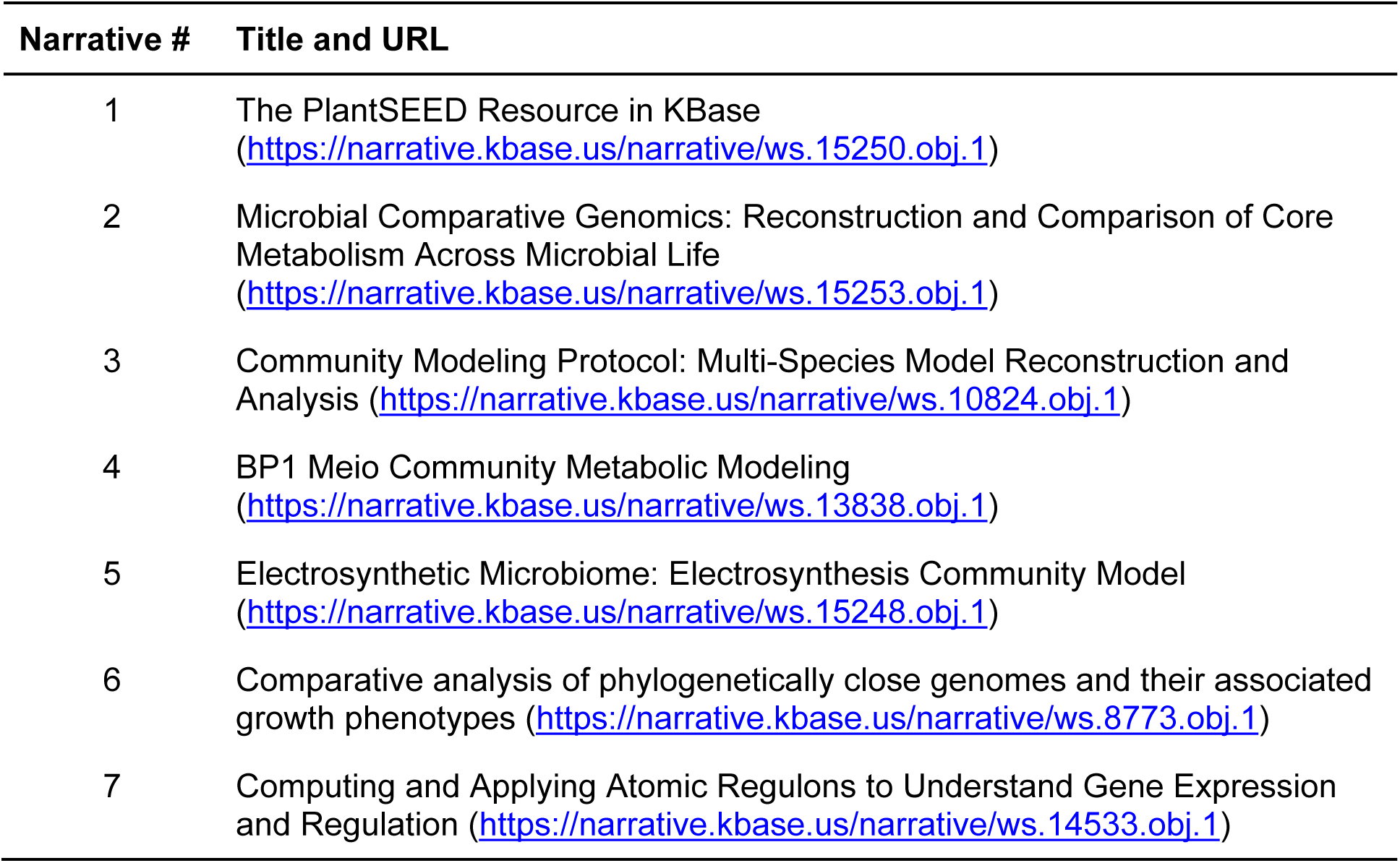
Example Narratives Demonstrating the Use of KBase (sign up for a KBase User Account at www.kbase.us before clicking the links to view these Narratives.)

KBase also has a sophisticated, unique pipeline for microbiome modeling and analysis, as demonstrated by three recent publications. One is a book chapter (www.kbase.us/community-modeling/) that explores various microbiome modeling paradigms, predicting potential interactions between the gut microbes *Bacteroides thetaiotaomicron* and *Faecalibacterium prausnitzii* as a case study (Table 2, Narrative 3)^16^. Another analysis examines interactions between an autotrophic carbon-fixing cyanobacteria, *Thermosynechococcus elongatus* BP-1, and the heterotrophic gram-positive species, *Meiothermus ruber* Strain A. Model predictions from this analysis were validated by a combination of published growth-condition data and comparison of model-predicted fluxes with metatranscriptome-based expression data (Table 2, Narrative 4)^28^. Finally, metagenomic and metatranscriptomic data from an electrosynthetic microbiome were assembled into genomes, models, and flux predictions for three dominant species in the microbiome (Table 2, Narrative 5)^29^. To our knowledge, no other platform is capable of performing the combination of community modeling analyses and expression data comparisons demonstrated in these featured Narratives.

KBase also has powerful comparative genomics tools, including newly developed apps for analyzing phenotype data. These tools are highlighted in a recent publication in which two phylogenetically close genomes are systematically compared, identifying how changes in gene content result in changes in growth phenotypes^30^ (Table 2, Narrative 6)^31^. In this analysis, an existing metabolic model^32^ is propagated to another strain of the same species. The models are then applied to understand the differences between Biolog data gathered for both species, including identifying growth conditions where additional experimental validation is needed. The models are also applied to predict essential and nonessential metabolic genes, with validation performed via comparison with a recently published TN-seq dataset^33^.

In another recent publication leveraging KBase’s range of expression tools, expression data are loaded for five genomes: *Escherichia coli, S. oneidensis, Pseudomonas aeruginosa, Thermus thermophilus,* and *Staphylococcus aureus* (Table 2, Narrative 7)^20^. KBase apps are used to identify clusters of co-expressed genes in these genomes, and the KBase Software Development Kit (SDK) is applied to add a new app for computing these clusters. This new app is compared to the others, and co-expressed clusters are also compared across all five genomes.

These peer-reviewed studies demonstrate KBase’s wide range of capabilities and the power of Narratives in disseminating complex computational analyses that are well documented, easily reproducible, and extensible.

### KBase Software Development Kit

Computational approaches to analyzing biological data are heterogeneous and can evolve rapidly. Accordingly, the KBase platform must be able to integrate new software tools from diverse sources with varying computational requirements while still maintaining a consistent data model with data provenance and version history. This integration is accomplished using the KBase SDK, a set of command-line tools and a web interface enabling any developer or advanced user to build, test, register, and deploy new or existing software as KBase apps. To facilitate transparent, reproducible, and open science, all software contributed to the central KBase software repository must adhere to a standard open-source license (https://opensource.org/licenses). Information about the app developer is maintained in the user documentation for that app so that credit can be properly given to the contributor. Data provenance, job management, usage logging, and app versioning are handled automatically by the platform, allowing developers to contribute new scientific tools quickly with minimal KBase-specific training.

Scientific computation in KBase is managed with a distributed Docker-based^34^ execution model. Docker allows the individual programs and system dependencies of each Narrative app to be saved as an image and run securely and identically across physical resources within individual, isolated environments called containers. The SDK enables developers to build and configure images and containers in a standard way, making them directly accessible to Narrative apps. The Docker execution model is powerful because contributors can test code on their own machine exactly as it runs on KBase production systems, and containers can be deployed and executed as soon as they are registered without any other manual intervention or system deployments. Docker images also provide a basis for reproducible execution of apps in Narratives.

KBase apps built with the SDK include a wide and growing collection of systems biology tools. The SDK has already been applied by KBase developers and external users to implement all of the tools presently available in KBase, including all tools mentioned in the “Alice and Bob” use case example and the science applications described above.

## Comparison with Other Platforms

KBase builds on many ideas from previous systems and infrastructures designed to support large-scale bioinformatics analysis and model building. Many KBase developers have also worked on other systems including the SEED^35^, MicrobesOnline^36^, RAST^37^, PATRIC^38^, MG-RAST^39^, ModelSEED^15^, Gramene^40^, iPlant (now CyVerse)^41^, RegTransBase^42^, RegPrecise^43^, and others.

KBase differs from existing systems in several ways. While platforms such as Galaxy^44^, Taverna^45^, CyVerse, XSEDE^46^, myExperiment^47^, and GenePattern^48^ permit a user to develop and run complex bioinformatics workflows, they currently lack tools for workflow annotation or predictive modeling. In contrast, systems such as COBRA Toolbox^49^, Pathway Tools^50^, and RAVEN Toolbox^51^ support predictive modeling of metabolism but lack support for genome assembly, annotation, and comparison. More general computational platforms like Kepler^52^, Pegasus^53^, Globus^54^, and Jupyter^5^ extensively support data and workflow management but lack integration with bioinformatics analysis tools. Most of these tools do not allow formal integration of user data with organized reference data or sharing models for data and analyses.

A key distinction between KBase and such systems is KBase’s extensible core data model, whose aim is to provide consistent access to and integration of reference data sets and user-contributed data across all applications, user interfaces, and services such as annotation from well-curated ontologies. These features mean that KBase users do not have to adjust data files or formats once data are loaded into the system and that both user and reference data can be analyzed using the same tools.

KBase serves many of the emerging requirements for open, verifiable science supported wholly or in part by systems such as Galaxy, CyVerse, and Synapse^55^. These capabilities include promoting reproducibility and transparency by persistently linking data and the analyses used to produce them. KBase also improves user accessibility to sophisticated computing by providing an environment populated with tools and data that users can apply on a high-end computing infrastructure without additional setup or maintenance of any software or hardware. Granular sharing with designated individuals or the community at large is supported as well, with increasing capabilities for collaborative analysis and editing of data, experiments, and results.

The sharing, collaborative editing, and emerging user-discovery systems in KBase are the first steps toward a social networking and scientific project management system enabling users to find other researchers working in their areas, form teams, organize data and analytical projects, and communicate their results effectively.

While some systems such as Synapse and Galaxy do provide provenance information on how particular data files were produced, KBase supports a rich provenance system that integrates such information with both the KBase data model and social network. This integration enables more effective discovery of newly integrated reference data and the relevant work products of other users, paving the way toward a system continually enriched by user contributions and interactions.

## Methods

### System Architecture

KBase has a scalable service-oriented architecture (Fig. 4) in which core components communicate through web APIs. The Narrative user interface is the central hub where users submit computational tasks and manage data. KBase’s “Execution Engine” uses a distributed execution model that queues and routes tasks to the appropriate physical compute resources of the internal cloud or HPC facilities. Scientific analysis software is exposed through Narrative apps built and dynamically registered with KBase using the SDK. The software made accessible by apps and any system dependencies is captured and versioned using Docker^34^. The App Catalog maintains the dynamic repository of apps and Docker images and serves this information to the user interface and Execution Engine on demand. Reference and user data are stored identically in a system composed of structured data objects, flat files, and structured references. The references define relationships among types of data objects, enabling a traversable, integrated data model. A set of data services maintains versioning, provenance, permissions, and searching.

**Figure 4.**
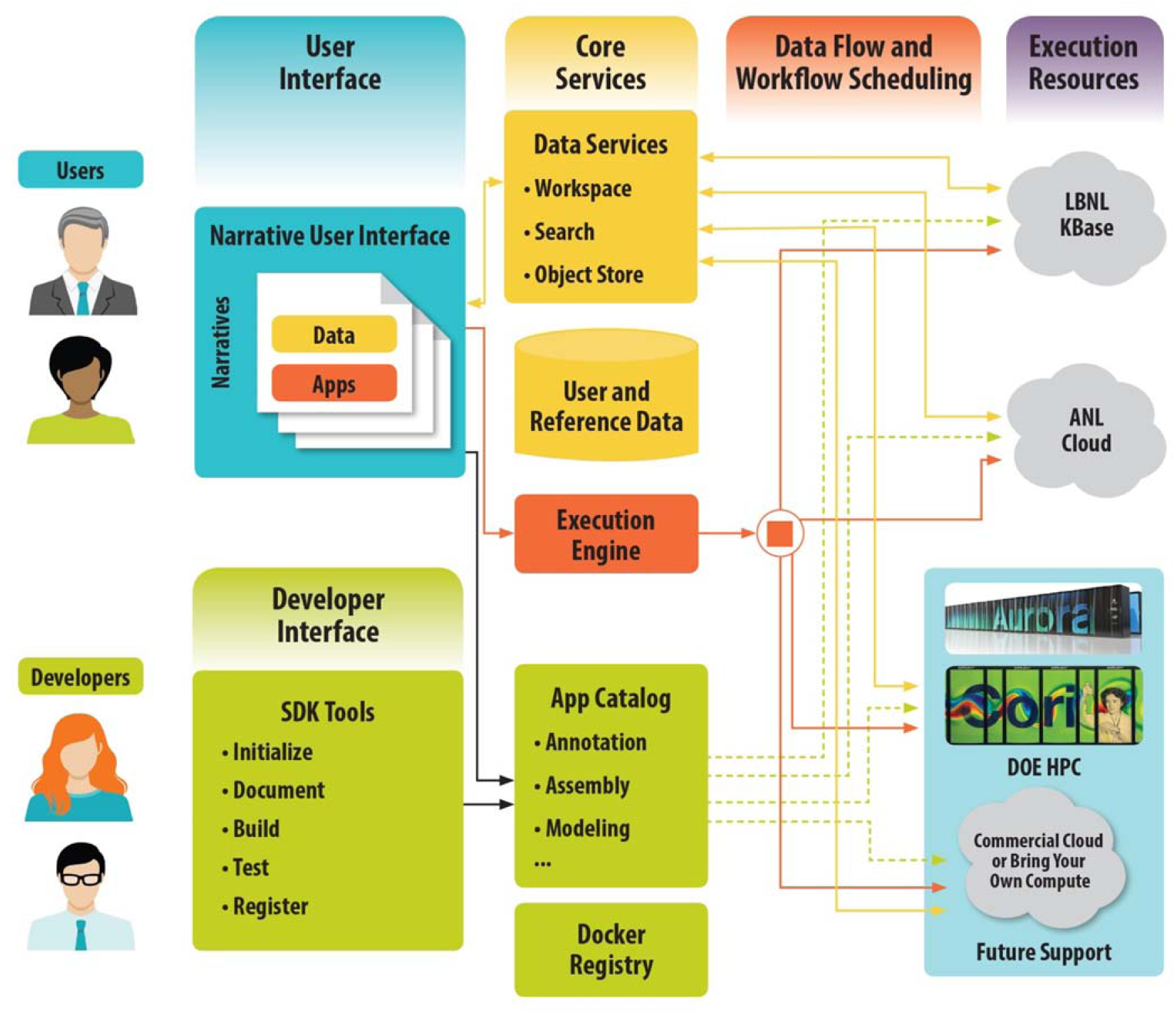
*KBase high-level architecture. KBase is based loosely on a service-oriented architecture that bundles related functionality into a set of independently scalable services that are managed to provide responsive interaction via the Narrative Interface. [Supercomputer images of Aurora courtesy Argonne National Laboratory (under a <u>Creative Commons license</u>) and Cori courtesy the DOE National Energy Research Scientific Computing Center (NERSC) run by Lawrence Berkeley National Laboratory.]*

The physical infrastructure hosting the KBase software platform operates at Argonne National Laboratory and Lawrence Berkeley National Laboratory using an OpenStack managed cloud in combination with large-scale computers at DOE’s National Energy Research Scientific Computing Center (NERSC) and Argonne Leadership Computing Facility (ALCF). Multiple deployments of platform services across sites provide some fault tolerance and failover capabilities.

## Code Availability

KBase software, available at https://github.com/kbase, is open source and freely distributed under the MIT License.

## Discussion

KBase has made some significant strides in leveraging the opportunities of this new data-rich era of biology. Important developments include (1) a detailed data model with support for provenance and versioning; (2) a rich and growing body of powerful analytical and modeling tools; (3) a collaborative user interface that seamlessly integrates data, analyses, and commentary to capture deep, reproducible scientific analyses; (4) an SDK enabling third-party developers to extend the system with new tools; and (5) a robust, scalable, and extensible underlying computational infrastructure.

Since its first public production release in February 2015, the KBase platform has been used extensively by hundreds of users who have built thousands of Narratives. Among these analyses is work described in over 10 peer-reviewed publications covering a wide range of topics. KBase is the only platform where users can immediately apply many of the analysis tools used in these publications in a turnkey, compute-, and data-ready environment using a graphical interface.

Extensive user testing is being done to improve the user experience offered by KBase, as well as the range of apps. User feedback indicates that the user interface, with its graphical access to data and apps, enables researchers to quickly learn how to run sophisticated multi-step analyses, find collaborators, and share results. The current KBase release also contains prototype functionality allowing researchers to more fully leverage the power of the Jupyter Notebook by using use code cells to programmatically access KBase and build custom analyses. As KBase improves support for its programming interface and lowers the barrier for third-party incorporation of apps and data types, we anticipate rapid growth in platform functionality and adoption by the community.

A key area of improvement in the short term is extending the number and variety of apps and associated data types contributed by KBase personnel and external developers. Development will focus on expanding metagenomics capabilities, improving support for eukaryotic genomic analyses, and providing more-comprehensive comparative genomics tools. Upcoming developments specifically will include improving the input and processing of bulk data sets, easing the process for defining new data types and their relationships to other information in the system, enabling large-scale execution on cloud and HPC resources, and adding frameworks that simplify creation of visualizations.

KBase also plans to extend the social platform and user interface to allow (1) more flexible discovery of users; (2) formation of “projects” organizing people, data, and Narratives that can control their group privacy and sharing options more effectively; and (3) a formal process for publishing Narratives.

## Acknowledgements

This work is supported by the Office of Biological and Environmental Research’s Genomic Science program within the U.S. Department of Energy Office of Science, under award numbers DE-AC02-05CH11231, DE-AC02-06CH11357, DE-AC05-00OR22725, and DE-AC02- 98CH10886.

## Contributions

APA, RLS, RWC, SM, CSH, PD, DW and FP developed the concept and vision.

APA, RLS, SC, MWS, MLH, WJR, DG, DMO, SYC, TSB, FM, DC, JB, AAB, BB, SEB, CCB, JMC, JC, RC, NC, JJD, MDJ, SD, FH, MPJ, KPK, BO, SP, EP, GAP, SR, PR, SMDS, MS, RAS, MHS, JT, FX, HSY and SY designed and developed the system.

RWC, NLH, RK, DC, DJW, EMG, BHD, SK, BHA, ED, MMD, ID, JNE, GF, JPF, PMF, WG, MG, JG, RJ, SNK, VK, MLL, MDM, PN, OTY, GJO, BDP, SSP, PCR, MCS, NLT and DFW developed, documented and conducted testing and validation.

APA and CSH drafted the manuscript.

NLH, HLH, BHA, MDM, MPJ, AAB, JMC, DC, BO, BHD, NLT, SM, PCR and MDJ revised the manuscript and provided important intellectual content.

APA, RWC and CSH reviewed and approved the final version to be published.

## References

1 Prlić, A. & Procter, J. B. Ten Simple Rules for the Open Development of Scientific Software. PLOS Computational Biology 8, e1002802, doi:10.1371/journal.pcbi.1002802 (2012).

2 Millman, K. J., and Fernando Pérez. in Implementing reproducible research (ed Friedrich Leisch Victoria Stodden, and Roger D. Peng) 149–183 (CRC Press, 2014).

3 Stodden, V. et al. Enhancing reproducibility for computational methods. Science 354, 1240–1241, doi:10.1126/science.aah6168 (2016).

4 DOE Systems Biology Knowledgebase Implementation Plan, http://genomicscience.energy.gov/compbio/kbaseplan/index.shtml (2010).

5 Perez, F. & Granger, B. E. IPython: A system for interactive scientific computing. Comput Sci Eng 9, 21–29, doi:10.1109/MCSE.2007.53 (2007).

6 O’Leary, N. A. et al. Reference sequence (RefSeq) database at NCBI: current status, taxonomic expansion, and functional annotation. Nucleic Acids Res 44, D733–745, doi:10.1093/nar/gkv1189 (2016).

7 Goodstein, D. M. et al. Phytozome: a comparative platform for green plant genomics. Nucleic Acids Res 40, D1178–1186, doi:10.1093/nar/gkr944 (2012).

8 Hubbard, T. et al. The Ensembl genome database project. Nucleic Acids Res 30, 38–41 (2002).

9 Ashburner, M. et al. Gene ontology: tool for the unification of biology. The Gene Ontology Consortium. Nat Genet 25, 25–29, doi:10.1038/75556 (2000).

10 Overbeek, R. et al. The subsystems approach to genome annotation and its use in the project to annotate 1000 genomes. Nucleic Acids Res 33, 5691–5702, doi:10.1093/nar/gki866 (2005).

11 Finn, R. D. et al. Pfam: the protein families database. Nucleic Acids Res 42, D222-230, doi:10.1093/nar/gkt1223 (2014).

12 Mulder, N. J. et al. New developments in the InterPro database. Nucleic Acids Res 35, D224–228 (2007).

13 Aziz, R. K. et al. The RAST Server: rapid annotations using subsystems technology. BMC Genomics 9, 75 (2008).

14 Price, M. N., Dehal, P. S. & Arkin, A. P. FastTree 2–approximately maximum-likelihood trees for large alignments. PLoS One 5, e9490, doi:10.1371/journal.pone.0009490 (2010).

15 Henry, C. S. et al. High-throughput generation, optimization and analysis of genomescale metabolic models. Nat Biotechnol 28, 977–982, doi:10.1038/nbt.1672 (2010).

16 Faria, J. P. et al. in Hydrocarbon and Lipid Microbiology Protocols 1–27 (Humana Press, 2016).

17 Latendresse, M. Efficiently gap-filling reaction networks. BMC Bioinformatics 15, 225, doi:10.1186/1471-2105-15-225 (2014).

18 Dreyfuss, J. M. et al. Reconstruction and validation of a genome-scale metabolic model for the filamentous fungus Neurospora crassa using FARM. PLoS Comput Biol 9, 096354, doi:10.1371/journal.pcbi.1003126 (2013).

19 Ghosh, S. & Chan, C. K. Analysis of RNA-Seq Data Using TopHat and Cufflinks. Methods Mol Biol 1374, 339–361, doi:10.1007/978-1-4939-3167-5_18 (2016).

20 Faria, J. P. et al. Computing and Applying Atomic Regulons to Understand Gene Expression and Regulation. Frontiers in Microbiology 7, doi:10.3389/fmicb.2016.01819 (2016).

21 Deutschbauer, A. et al. Evidence-based annotation of gene function in Shewanella oneidensis MR-1 using genome-wide fitness profiling across 121 conditions. PLoS Genet 7, e1002385, doi:10.1371/journal.pgen.1002385 (2011).

22 Ong, W. K. et al. Comparisons of Shewanella strains based on genome annotations, modeling, and experiments. BMC Syst Biol 8, 31, doi:10.1186/1752-0509-8-31 (2014).

23 Copeland, A. et al. Complete sequence of Shewanella amazonensis SB2B, EMBL/GenBank/DDBJ databases https://www.ncbi.nlm.nih.gov/genome/1223 (2006).

24 Venkateswaran, K., Dollhopf, M. E., Aller, R., Stackebrandt, E. & Nealson, K. H. Shewanella amazonensis sp. nov., a novel metal-reducing facultative anaerobe from Amazonian shelf muds. Int J Syst Bacteriol 48, 965–972, doi:10.1099/00207713-48-3-965 (1998).

25 Wetmore, K. M. et al. Rapid quantification of mutant fitness in diverse bacteria by sequencing randomly bar-coded transposons. MBio 6, e00306-00315, doi:10.1128/mBio.00306-15 (2015).

26 Seaver, S. M. et al. High-throughput comparison, functional annotation, and metabolic modeling of plant genomes using the PlantSEED resource. Proceedings of the National Academy of Sciences of the United States of America 111, 9645–9650, doi:10.1073/pnas.1401329111 (2014).

27 Edirisinghe, J. N. et al. Modeling central metabolism and energy biosynthesis across microbial life. BMC Genomics 17, 568, doi:10.1186/s12864-016-2887-8 (2016).

28 Henry, C. S. et al. Microbial Community Metabolic Modeling: A Community Data-Driven Network Reconstruction. J Cell Physiol 231, 2339–2345, doi:10.1002/jcp.25428 (2016).

29 Marshall, C. et al. Electron transfer and carbon metabolism in an electrosynthetic microbial community, Preprint at http://biorxiv.org/content/early/2016/07/07/059410 (2016).

30 Broberg, C. A., Wu, W., Cavalcoli, J. D., Miller, V. L. & Bachman, M. A. Complete Genome Sequence of Klebsiella pneumoniae Strain ATCC 43816 KPPR1, a RifampinResistant Mutant Commonly Used in Animal, Genetic, and Molecular Biology Studies. Genome Announc 2, doi:10.1128/genomeA.00924-14 (2014).

31 Henry, C. S. et al. Generation and validation of the iKp1289 metabolic model for Klebsiella pneumoniae KPPR1. Journal of Infectious Disease In press (2016).

32 Liao, Y. C. et al. An experimentally validated genome-scale metabolic reconstruction of Klebsiella pneumoniae MGH 78578, iYL1228. J Bacteriol 193, 1710–1717, doi:10.1128/JB.01218-10 (2011).

33 Bachman, M. A. et al. Genome-Wide Identification of Klebsiella pneumoniae Fitness Genes during Lung Infection. MBio 6, e00775, doi:10.1128/mBio.00775-15 (2015).

34 Anderson, C. Docker. IEEE Software 32, 102–105 (2015).

35 Overbeek, R., Disz, T. & Stevens, R. The SEED: A peer-to-peer environment for genome annotation. Commun ACM 47, 46–51, doi:Doi 10.1145/1029496.1029525 (2004).

36 Dehal, P. S. et al. MicrobesOnline: an integrated portal for comparative and functional genomics. Nucleic Acids Res 38, D396–400, doi:10.1093/nar/gkp919 (2010).

37 Brettin, T. et al. RASTtk: a modular and extensible implementation of the RAST algorithm for building custom annotation pipelines and annotating batches of genomes. Sci Rep 5, 8365, doi:10.1038/srep08365 (2015).

38 Wattam, A. R. et al. PATRIC, the bacterial bioinformatics database and analysis resource. Nucleic Acids Res 42, D581–591, doi:10.1093/nar/gkt1099 (2014).

39 Meyer, F. et al. The metagenomics RAST server - a public resource for the automatic phylogenetic and functional analysis of metagenomes. BMC bioinformatics 9, 386, doi:10.1186/1471-2105-9-386 (2008).

40 Ware, D. et al. Gramene: a resource for comparative grass genomics. Nucleic Acids Res 30, 103–105 (2002).

41 Goff, S. A. et al. The iPlant Collaborative: Cyberinfrastructure for Plant Biology. Frontiers in plant science 2, 34, doi:10.3389/fpls.2011.00034 (2011).

42 Cipriano, M. J. et al. RegTransBase–a database of regulatory sequences and interactions based on literature: a resource for investigating transcriptional regulation in prokaryotes. BMC Genomics 14, 213, doi:10.1186/1471-2164-14-213 (2013).

43 Novichkov, P. S. et al. RegPrecise 3.0–a resource for genome-scale exploration of transcriptional regulation in bacteria. BMC Genomics 14, 745, doi:10.1186/1471-2164- 14–745 (2013).

44 Goecks, J., Nekrutenko, A., Taylor, J. & Galaxy, T. Galaxy: a comprehensive approach for supporting accessible, reproducible, and transparent computational research in the life sciences. Genome Biol 11, R86, doi:10.1186/gb-2010-11-8-r86 (2010).

45 Oinn, T. et al. Taverna: a tool for the composition and enactment of bioinformatics workflows. Bioinformatics 20, 3045–3054, doi:10.1093/bioinformatics/bth361 (2004).

46 Towns, J. et al. XSEDE: Accelerating Scientific Discovery. Comput Sci Eng 16, 62–74 (2014).

47 Goble, C. A. et al. myExperiment: a repository and social network for the sharing of bioinformatics workflows. Nucleic Acids Res 38, W677–682, doi:10.1093/nar/gkq429 (2010).

48 Reich, M. et al. GenePattern 2.0. Nat Genet 38, 500–501, doi:10.1038/ng0506-500 (2006).

49 Schellenberger, J. et al. Quantitative prediction of cellular metabolism with constraint-based models: the COBRA Toolbox v2.0. Nat Protoc 6, 1290–1307, doi:10.1038/nprot.2011.308 (2011).

50 Karp, P. D. et al. The EcoCyc and MetaCyc databases. Nucleic Acids Res 28, 56–59 (2000).

51 Agren, R. et al. The RAVEN toolbox and its use for generating a genome-scale metabolic model for Penicillium chrysogenum. PLoS Comput Biol 9, e1002980, doi:10.1371/journal.pcbi.1002980 (2013).

52 Altintas, I. et al. Kepler: An extensible system for design and execution of scientific workflows. 16th International Conference on Scientific and Statistical Database Management, Proceedings, 423–424 (2004).

53 E. D. Pegasus: A framework for mapping complex scientific workflows onto distributed systems. Sci Programming-Neth 13, 219–237 (2005).

54 Ananthakrishnan, R., Chard, K., Foster, I. & Tuecke, S. Globus Platform-as-a-Service for Collaborative Science Applications. Concurr Comp-Pract E 27, 290–305, doi:10.1002/cpe.3262 (2015).

55 Derry, J. M. J. et al. Developing predictive molecular maps of human disease through community-based modeling. Nature Genetics 44, 127–130, doi:10.1038/ng.1089 (2012).

